# Altering brain dynamics with transcranial random noise stimulation

**DOI:** 10.1101/227140

**Authors:** Onno van der Groen, Nicole Wenderoth, Jason B. Mattingley

## Abstract

Random noise can enhance the detectability of weak signals in nonlinear threshold systems, a phenomenon known as stochastic resonance (SR). This concept is not only applicable to single threshold systems but can also be applied to dynamical systems with multiple attractor states, such as observed during the phenomenon of binocular rivalry. Binocular rivalry can be characterized by marginally stable attractor states between which the brain switches in a spontaneous, stochastic manner. The switches are thought to be driven by a combination of neuronal adaptation and noise. Here we used a computational model to predict the effect of noise on perceptual dominance durations when either low-contrast or high-contrast stimuli are presented. Subsequently we compared the model prediction to a series of three experiments where we measured binocular rivalry dynamics when noise (zero-mean Gaussian white noise) was added either to the visual stimulus (Exp. 1) or directly to the visual cortex (Exp. 2 and Exp. 3) by applying transcranial Random Noise Stimulation (tRNS 1mA, 100-640 Hz zero mean Gaussian white noise). We found that adding noise significantly reduced the mixed percept duration (Exp. 1 and Exp. 2). This effect was only present for low-contrast but not for high-contrast visual stimuli which is in line with the model predictions. Our results demonstrate that both central and peripheral noise can influence state-switching dynamics of binocular rivalry under specific conditions (e.g. low visual contrast stimuli), in line with a SR-mechanism.

## Introduction

Noise is detrimental for the transfer of information in linear systems (McDonnell & Abbott, 2009). However, in nonlinear systems such as the brain, noise can enhance information transfer via a stochastic resonance (SR) mechanism (McDonnell & Abbott, 2009; Ward, Neiman, & Moss, 2002). SR can be experimentally observed when an optimal level of noise is added to a non-linear system which enhances (i) the output of the system e.g. by improving the signal-to-noise ratio (SNR) (Russell, Wilkens, & Moss, 1999; van der Groen & Wenderoth, 2016), (ii) the signal amplitude (Manjarrez, Diez-Martinez, Mendez, & Flores, 2002) or (iii) the degree of coherence within neural networks (Ward, MacLean, & Kirschner, 2010). In humans, the SR-effect has been observed in multiple sensory modalities when both signal and experimentally controlled noise are added to the peripheral nervous system (Collins, Imhoff, & Grigg, 1996, 1997; Simonotto et al., 1997; Zeng, Fu, & Morse, 2000). Recently, we have demonstrated that central mechanisms of perception are also sensitive to a SR-effect (van der Groen & Wenderoth, 2016). We showed that an optimal level of transcranial random noise stimulation (tRNS), a form of non-invasive brain stimulation, applied over visual cortex can enhance the detection performance of weak subthreshold visual stimuli.

Theoretical considerations predict that SR does not only play a role in signal enhancement but that it can also influence the dynamics of bistable-systems (L. Gammaitoni, Marchesoni, Menichella-Saetta, & Santucci, 1989). One paradigm that allows the observation and measurement of how the brain dynamically transitions between different perceptual states is binocular rivalry. Binocular rivalry is a perceptual phenomenon that occurs when different stimuli are simultaneously presented to each eye. During binocular rivalry visual awareness switches spontaneously between the two stimuli (Levelt, 1965) so that at any given time participants perceive either one of the two images (exclusive percept) or a combination of both (mixed percept).

Models of binocular rivalry propose that it reflects a competition between neural populations coding for each image (Tong, Meng, & Blake, 2006). The neural population coding for the dominant percept inhibits neurons that code for the suppressed image. However, over time the inhibition of the dominant population becomes weaker due to adaptation or fatigue, which allows the suppressed image to become dominant (Blake, 1989; Tong et al., 2006). This results in a deterministic anti-phase oscillation of the firing rates of the two neuronal populations (Shpiro, Moreno-Bote, Rubin, & Rinzel, 2009). However, if adaptation was the only driving factor of binocular rivalry, perception would change fairly regularly.

In fact, the dynamics of binocular rivalry are highly nonlinear and stochastic, leading to the proposal that noise associated with the activity of the two neuronal populations causes a random distribution of dominance durations (Lankheet, 2006). Accordingly, noise is thought to play a crucial role in the occurrence of the perceptual switches and it has been suggested to represent an essential driving force of rivalry dynamics (Brascamp, van Ee, Noest, Jacobs, & van den Berg, 2006; Huguet, Rinzel, & Hupe, 2014). Consequently, there are many successful models of binocular rivalry which contain a noise component to describe the rivalry dynamics (Moreno-Bote, Rinzel, & Rubin, 2007; Said & Heeger, 2013; Shpiro et al., 2009). While it has been demonstrated that rivalry dynamics can be influenced by adaptation (Roumani & Moutoussis, 2012), the effect of adding noise directly to the brain during binocular rivalry has not been empirically tested. Rivalry dynamics can conceptually be represented by a double-well energy landscape, with each well representing one of the images (see fig 1). Neuronal adaptation can be represented as changes in the energy-landscape, that is, adaptation will reduce the depth of the well indicating that the current state becomes less stable, thus, making a switch to the competing precept more likely.

**Figure 1:**
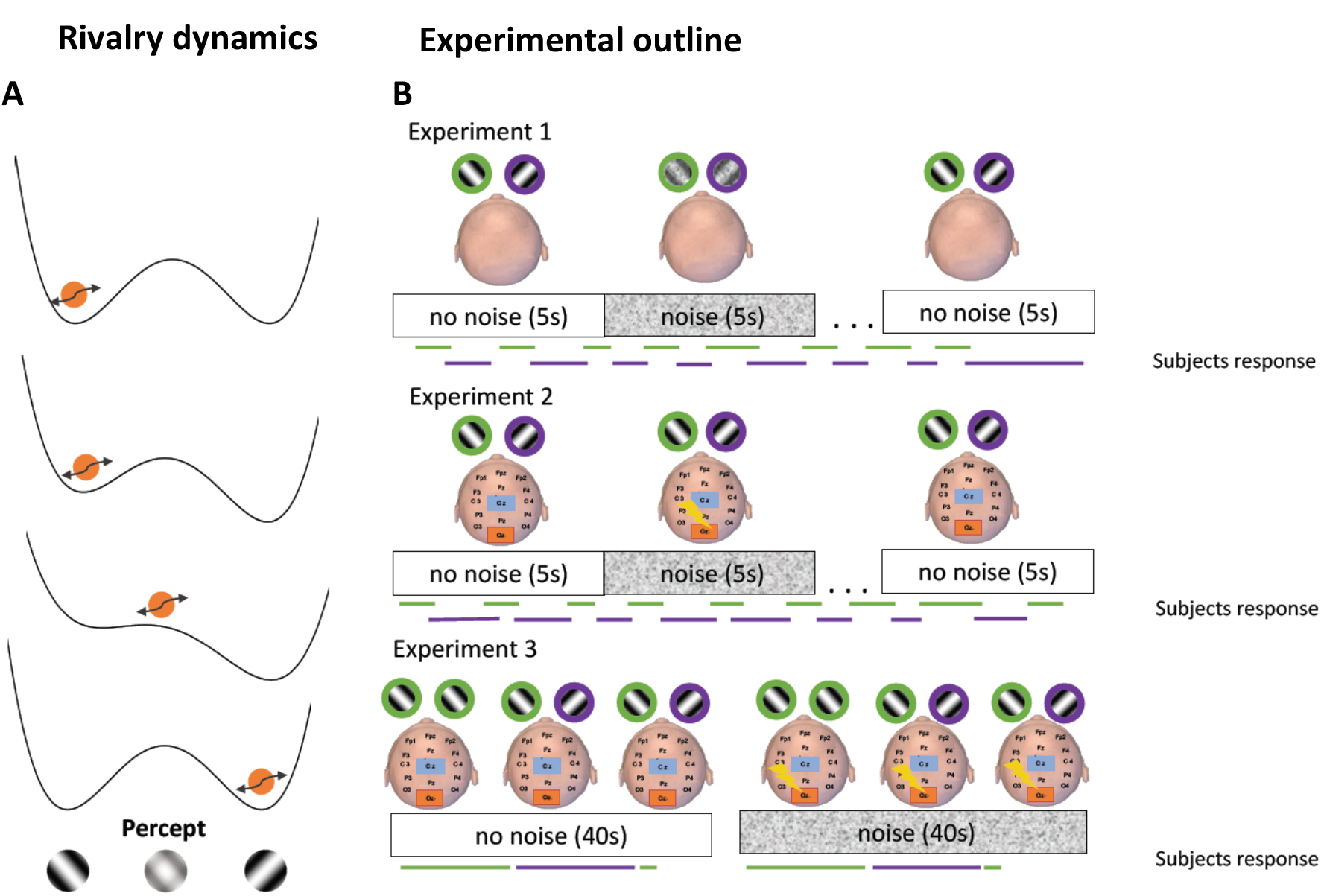
**A)** Representation of the rivalry dynamics. Binocular rivalry can be represented by a double-well energy landscape. The orange ball determines the current percept. Binocular rivalry is thought to be driven by adaptation and noise. Adaptation changes the landscape, meaning one of the wells becomes less shallow. Noise (arrow) causes the percept to change more quickly when the boundary between the two wells is low. **B)** Experimental outline. In experiment 1 noise (zero-mean Gaussian white noise) was applied to the visual stimulus for 5 seconds followed by 7 seconds of no stimulation. The same noise was applied to the left and right eye. The noise intensity was subthreshold for each individual participant. Experiment 2 followed the same protocol, except that the noise was applied to the visual cortex directly with tRNS (zero-mean Gaussian white noise, 100–640 Hz). In experiment 3 noise was also applied to the visual cortex with tRNS. However, two images with the same orientation were presented to each eye at the start of each trial. After a variable interval one of the two images changed orientation, causing the percept to switch. Noise was applied in blocks of 40 seconds.

Counterintuitively, adding an optimal level of noise to such a system can change its dynamics (L. Gammaitoni et al., 1989; Rajasekar & Sanjuan). This can reduce the time spend in a single state by enhancing the strength of the weak driving signal (Rajasekar & Sanjuan). At a behavioural level this results in a decreased variability in time spent in a single state due to the oscillatory nature of the noise-enhanced driving signal, which is reflected in the model as a reduction of the standard deviation of the dominance durations. When the dynamics are only driven by the weak signal then we expect a low level of variation in the dominance durations. Here we investigate if the attractor dynamics of binocular rivalry can be modulated by adding noise to either the visual stimulus or directly to the visual cortex with tRNS (Onorato et al., 2016; Terney, Chaieb, Moliadze, Antal, & Paulus, 2008). Three experiments were performed: In experiment 1 noise was added to the visual stimulus to test if a SR-effect is induced when noise is added to the periphery. In experiment 2 we added noise to the visual cortex with tRNS to test if central mechanisms of perception are sensitive to a SR-effect. The results of these experiments suggest that rivalry dynamics can be influenced by noise, when there are three stable states. In experiment 3 we again added the noise to the visual cortex with tRNS, however, the experimental design was adapted in order to get only two stable states. In order to make clear predictions before data collection we simulated the effect of adding noise to rivalry dynamics with a computational model (Said & Heeger, 2013). We also modelled current flow in the brain (Thielscher, Antunes, & Saturnino, 2015) to estimate electric field strength in our region of interest (Truong et al., 2014).

## Materials and methods

The study was approved by the Kantonale Ethikkomission Zürich, Switzerland (KEK-ZH-Nr. 2014-0269) and by The University of Queensland Human Research Ethics Committee. Informed consent was obtained from all participants before the start of the experiment.

### General procedures of experiments 1 and 2

The data for experiments 1 and 2 were collected in Zurich. All experiments took place in a dark and quiet room. Visual stimuli (left and right tilted gratings) were generated using MATLAB version 2012b (MathWorks, Natick, USA) and the Psychophysics Toolbox (Brainard, 1997; Kleiner, Brainard, & Pelli, 2007; Pelli, 1997). Stimuli were controlled by a HP Elitedesk 800 G1 running Windows 7 (2009). Stimuli were presented on a Sony CPD-G420 color monitor with a calibrated linearized output at a resolution of 1280x1024 pixels, with a refresh rate of 75 Hz. The two images were placed on the horizontal meridian to the left and right hand side of the screen on a uniform grey background (53 Cd/m^2^). The visual stimuli were orthogonal Gaussian gratings (orientation ± 45°, diameter 3 cm, contrast 20 %, visual angle 4°, spatial-frequency 2.9 cycles per cm) surrounded by a white square (diameter 3.5 cm), to promote binocular fusion. Participants were seated in front of the monitor and viewed the images through a mirror stereoscope from a distance of 45 cm while resting their head on a chin rest. Each participant performed 8 runs of binocular rivalry, each lasting 3 minutes. The task for the participants was to continuously report on a keyboard whether they perceived the left or right tilted grating or a mixture of both. In each experiment the noise was applied in 4 runs to either the screen (exp. 1) or directly to the visual cortex with tRNS (exp. 2). The order of the noise runs was randomized over participants. In these runs the noise was applied for 5 seconds followed by a 5.5 – 7 seconds’ interval of no stimulation, and in total 18 times per run. In each experiment participants received a total of 360 seconds’ visual noise (exp. 1) or tRNS (exp. 2). Participants conducted one practice run without any noise before the start of the experiment.

### Experiment 1: Peri-noise condition

In this experiment, we tested if adding dynamic noise (zero-mean Gaussian white noise) to the visual stimuli influences binocular rivalry dynamics. The same noise was applied to the left and right eye. Before the start of the experiment we determined a rough-estimate of each individual’s noise threshold. Previous research demonstrated that a noise intensity corresponding to 60% of threshold effectively induces a SR-effect (van der Groen & Wenderoth, 2016). Therefore, we used a simple up-down method to estimate each individual’s 60% correct noise-threshold before the experiment started. We tested two cohorts of participant, one with a low contrast visual stimulus (20% contrast, n = 10, mean age = 23) and one with a high contrast visual stimulus (70% contrast, n = 10, mean age =24). It is well established that SR only occurs for weak stimuli, therefore, we expect to observe an SR effect only for the low contrast stimuli (Collins et al., 1996, 1997; Luca Gammaitoni et al., 1998; Simonotto et al., 1997; van der Groen & Wenderoth, 2016; Zeng et al., 2000).

### Experiment 2: tRNS-V1 condition

In this experiment, we tested the hypothesis that adding noise directly to the visual cortex with tRNS can influence rivalry dynamics. Noise was applied centrally with tRNS (100-640 Hz, zero-mean Gaussian white noise). Electrode placement was determined using the 10-20 system. The stimulation electrode was placed over the occipital region (Oz in the 10-20 EEG system) and the reference electrode over the vertex (Cz in the 10-20 EEG system). This setup has been demonstrated to be suitable for stimulation of the visual cortex (Neuling, Wagner, Wolters, Zaehle, & Herrmann, 2012). Electroconductive gel was applied to the contact side of the electrode (5x7 cm) to reduce skin impedance. Electrodes were held in place with a bandage. Stimulation was delivered by a battery-driven electrical stimulator (DC-Stimulator Plus, neuroConn). An intensity of 1 mA was applied since it has been demonstrated that this intensity effectively induces a SR-effect in most subjects (van der Groen & Wenderoth, 2016). The maximum current density in this experiment was 28.57 μA/cm^2^, which is within current safety limits (Fertonani, Ferrari, & Miniussi, 2015). We again tested two cohorts of participant, one where the visual stimulus had a low contrast (20% contrast, n = 15, mean age = 23) and one where the visual stimulus had a high contrast (70% contrast, n = 15, mean age = 24).

### Experiment 3: tRNS-V1 follow-up with optimized design

Data collection for this experiment took place at the Queensland Brain Institute in Brisbane, Australia. The visual stimuli in experiment 3 were the same as in experiments 1 and 2. Visual stimuli were generated using MATLAB version 2015a (MathWorks, Natick, USA) and the Psychophysics Toolbox (Brainard, 1997; Kleiner et al., 2007; Pelli, 1997). Stimuli were controlled by a Dell Precision T1700 running Windows 7 (2009). Stimuli were presented on an ASUS VG 248 QE colour monitor with a calibrated linearized output at a resolution of 1280x1024 pixels, with a refresh rate of 75 Hz. Participants (n = 15, mean age = 22) performed 8 runs, each made up of 6 x 40 second blocks. Each trial started with the two gratings presented with the same orientation in each eye (see fig. 1). After a variable interval (1-1.5 seconds) one of the gratings changed orientation, which results in this grating becoming the dominant percept. Over time, the percept automatically switches back to the orientation of the non-flipped grating, like during usual binocular rivalry. Therefore, in each trial two perceptual switches occurred. The first perceptual switch occurred after one of the gratings was flipped, the second perceptual switch occurred when the flipped grating lost dominance and the non-flipped grating became dominant. The amount of time the flipped grating was perceived is the dominance duration of the percept. The order in which one of the two gratings flipped was counterbalanced within each block. The task for the participants was to press the spacebar with their right hand as soon as their percept changed. After two spacebar presses or when the spacebar wasn’t pressed within 10 seconds of the first flip the next trial would start. Trials where the two perceptual switches didn’t occur were excluded from analysis. Participants performed as many trials as possible within a 40 second block. In half of the blocks, tRNS was applied to the visual cortex (same electrode setup as in exp. 2) for the duration of the block. A block with tRNS-stimulation was always followed by a block without stimulation in order to reduce any possible tRNS after effects. The order of the blocks with and without tRNS were counterbalanced over participants. In 4 runs the visual stimuli had a high contrast (70%), and in 4 runs a low contrast (20%). The order of grating contrast presentation was randomized over participants.

### Data analysis and statistics

Statistical analyses were performed using SPSS (version 20.0, IBM). The same statistical procedures were applied to all 3 experiments. For each participant, we calculated the mean dominance duration for the exclusive percepts. In experiment 1 and 2 we also calculated the mixed percept dominance durations. Times where no button was pressed, or when the dominance duration was shorter than 150 ms were excluded from analysis. Dominance durations terminated by the end of a block were not included in the analyses. In each experiment, we also calculated the number of perceptual switches. To test for the effect of the added noise, dominance durations, standard deviations and number of perceptual switches were subjected to a two-sided within-subject t-test. The a-level was set to 0.05 for all tests.

### Computational model of rivalry dynamics

We applied a computational model to predict how noise influences the rivalry dynamics (Said & Heeger, 2013). We used a conventional binocular rivalry model which relies on competition between neurons tuned to orthogonal orientations. The model relies on mutual inhibition and it also includes a noise component. The model contains two different neuronal populations and calculates the difference in firing rate between these populations. Over time the inhibition on the suppressed neuronal population weakens, which eventually results in the inhibited population becoming dominant and supressing the other population. When one population is firing fully the other population will be inhibited, resulting in the percept related to the fully firing population being dominant. When the difference in firing rate between the two populations is small then there is no winning population, resulting in a mixed percept. We introduced a criterion for mixed percepts, which was a difference in firing rate between the two neuronal populations of smaller than 0.1. In calculating mean dominance durations, we only included dominance times longer than 150 ms. In the model, we changed the strength of the visual stimulus and the amount of noise according to our experimental parameters. All other model parameters were identical to the original model parameters (Said & Heeger, 2013). The simulation was run 100 times in order to get an estimate of the variability of the model.

*Modelling of the electric field induced by tRNS* Modelling was used to estimate the electrical field strength in the visual cortex (Spheres 2.0 (Truong et al., 2014)). We did this for specific frequencies between 100 and 500 Hz (50 Hz steps, see table 1). We also modelled the current flow in the brain using the SIMNIBS toolbox (Thielscher et al., 2015). The modelling results show that the current reaches the brain and that it is focused on the visual cortex (figure 4). Our modelled electrical field strengths (table 1) show that all frequencies reach the cortex (scaling quasi-linearly with tRNS intensity).

**Table 1.** Modelled electric field strength (V/m) for different transcranial electrical current stimulation (tECSs) intensities (mA) and frequencies (Hz)

| *Frequency (HZ)* | *Intensity(mA)* |
| --- | --- |
|  | *1 mA* |
| 100 Hz | 2.2 V/m |
| 150 Hz | 2 V/m |
| 200 Hz | 2.2 V/m |
| 250 Hz | 2.2 V/m |
| 300 Hz | 2.1 V/m |
| 350 HZ | 2.1 V/m |
| 400 Hz | 2.1 V/m |
| 450 Hz | 2.2 V/m |
| 500 Hz | 2.2 V/m |

## Results

### Computational modelling results

Adding noise during binocular rivalry increases the duration of exclusive percepts (increase of 22% - average across low and high contrasts). The strongest effect of adding noise was observed as a substantial reduction of the mixed percept (63% decrease – averaged across low and high contrasts). This magnitude of this effect was different depending on contrast intensity: there was a stronger reduction of mixed percept durations for low contrast trials (72% change) than for high contrast trials (53% change).

### Behavioural results

#### Experiment 1

Adding noise to the low contrast visual stimulus during binocular rivalry significantly reduced the mean dominance duration of the mixed percept (t(9) = 2.712, p = 0.024, fig. 3). This effect coincided with a significant reduction in the standard deviation of the mixed percept (t(9) = 3, p = 0.015). Adding noise to a high contrast visual stimulus did not affect the mixed dominance duration (t(9) = 0.562, p = 0.588) or its standard deviation (t(9) = 0.234, p = 0.82).

**Figure 2:**
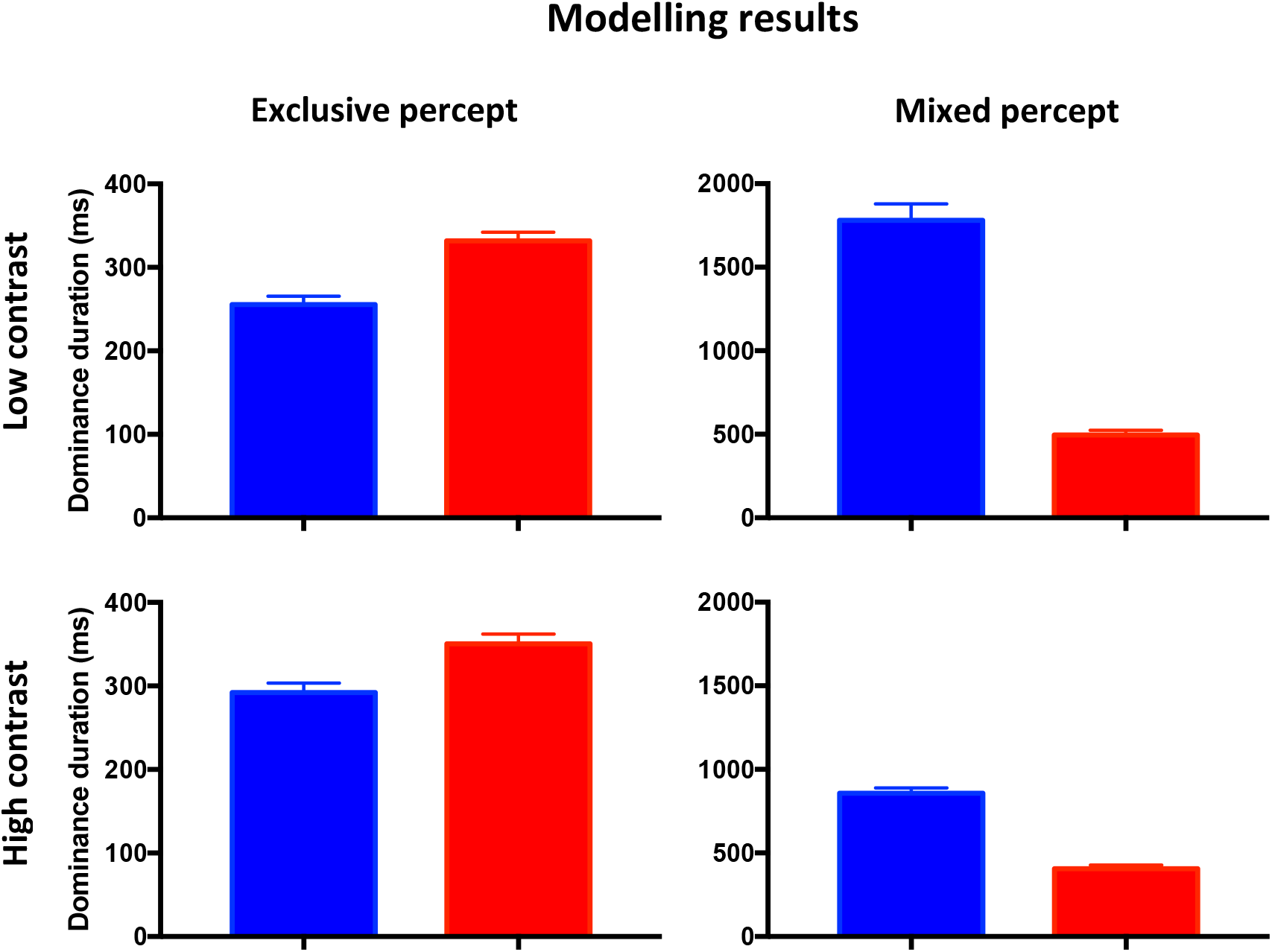
Computational modelling results. We modelled the effect of adding noise to a rivalry model, and determined dominance durations of the exclusive and mixed percepts (Said & Heeger, 2013). These results indicate that adding noise to the rivalry process mainly reduces the duration of the mixed percept for low contrast visual stimuli. For the exclusive percept duration, the dominance duration increases with an increasing noise level for both the low and high contrast visual stimuli.

**Figure 3:**
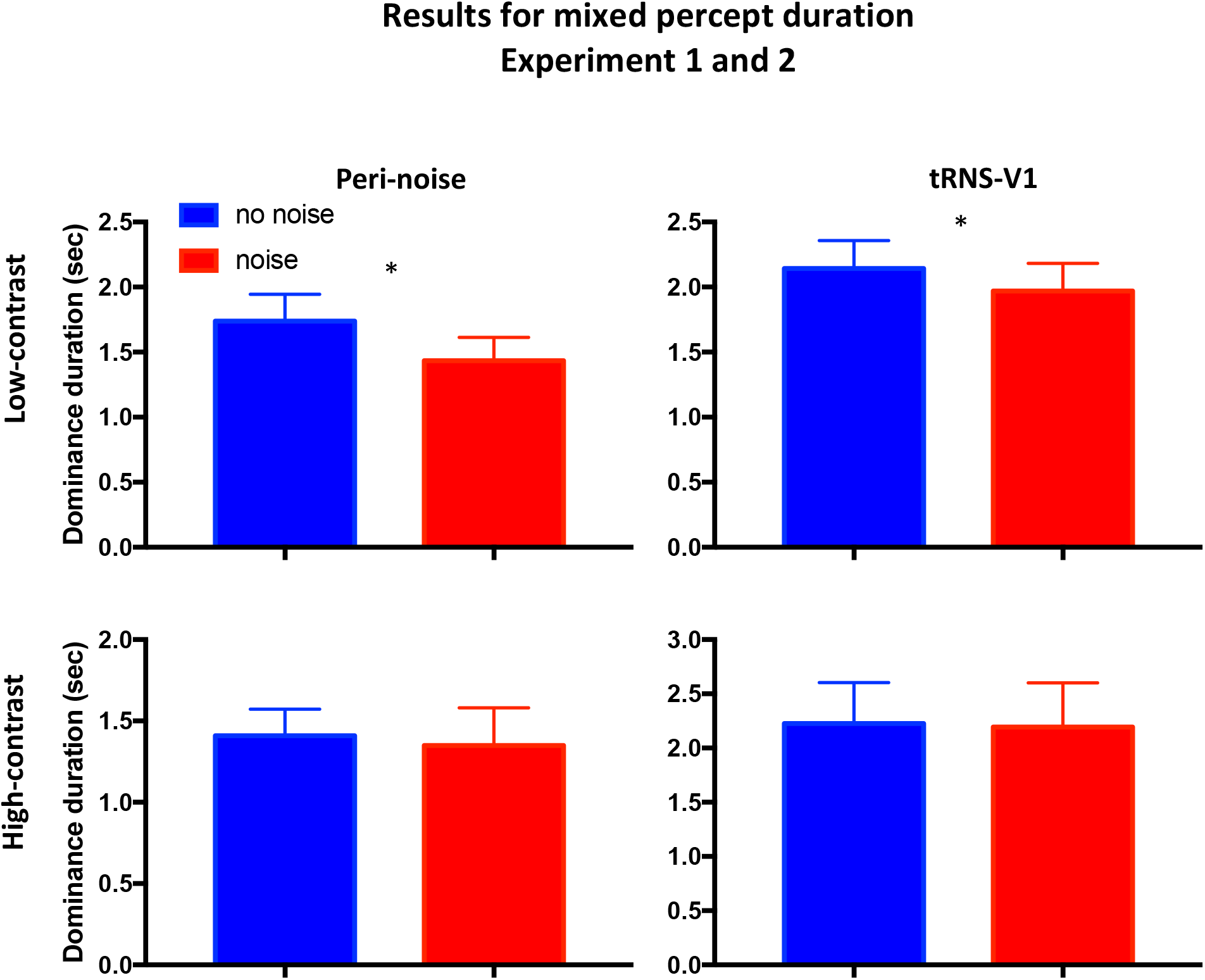
Behavioural results of experiments 1 (left) and 2 (right). Adding noise significantly reduces the dominance duration of the mixed percept for low contrast visual stimuli. There is no significant effect of adding noise on the dominance duration of the exclusive percept. Error bars represent SEM. *p < 0.05

The exclusive percepts were not significantly affected by peripheral noise for low contrast stimuli (t(9) = 0.789, p = 0.45) or high contrast stimuli (t(9) = 0.335, p = 0.75). These results are in line with the effect predicted by the computational model. The number of perceptual switches was not affected by noise (low contrast: t(9) = 0.65, p = 0.53; high contrast: t(9) = 0.75, p = 0.47).

#### Experiment 2

In experiment 2 we added noise directly to the visual cortex with tRNS. Consistent with previous investigations, no participant reported awareness of the tRNS stimulation during debriefing (Ambrus, Paulus, & Antal, 2010; Fertonani, Pirulli, & Miniussi, 2011).

Adding noise to visual cortex by tRNS yielded a similar pattern of results to experiment 1. That is, the mean mixed dominance duration of the low contrast visual stimulus was significantly reduced (t(14) = 2.581, p = 0.022, fig. 3). This effect coincided with a significant reduction in the standard deviation of the mixed percept (t(14) = 2.49, p = 0.026). Adding noise to a high contrast visual stimulus did not affect the mixed dominance duration (t(14) = 1.377, p = 0.19) or its standard deviation (t(14) = 1.129, p = 0.278). The exclusive percepts were not significantly affected by tRNS for the low contrast visual stimuli (t(14) = 0.832, p = 0.420) or for the high-contrast visual stimuli (t(14) = 1.044, p = 0.314). The number of perceptual switches were not affected by tRNS (low contrast: t(14) = 0.72, p = 0.485; high contrast: t(14) = 0.813, p = 0.43).

#### Experiment 3

It has been suggested that adding an optimal level of noise can influence the dynamics of bistable systems (Luca Gammaitoni et al., 1998). However, in experiments 1 and 2 the system had three marginally-stable states due to the mixed percept state. Therefore, we ran a follow-up experiment with a design which only allowed for two stable states in a different cohort of participants. We had to exclude two participants because they were not able to do the task. Our results did not reveal any significant effect of noise on the exclusive percept duration (low contrast: t(12) = 0.676, p = 0.51; high contrast: t(12) = 0.19, p = 0.85) or the number of perceptual switches (low contrast: t(12) = 1.26, p = 0.23; high contrast: t(12) = 1.01, p = 0.33). Taken together, our results suggest that adding noise to binocular rivalry preferentially effects escapes from the mixed percepts while it does not seem to affect exclusive percepts.

### tRNS electric field strength and current flow modelling

The modelling results show that that all stimulation frequencies were transmitted to the brain and that the applied current applied is sufficiently strong to reach the cortex (scaling quasi-linearly with tRNS intensity). The current is mainly focused on the visual cortex, however, there is spread to other brain areas (table 1 and fig. 4).

**Figure 4.**
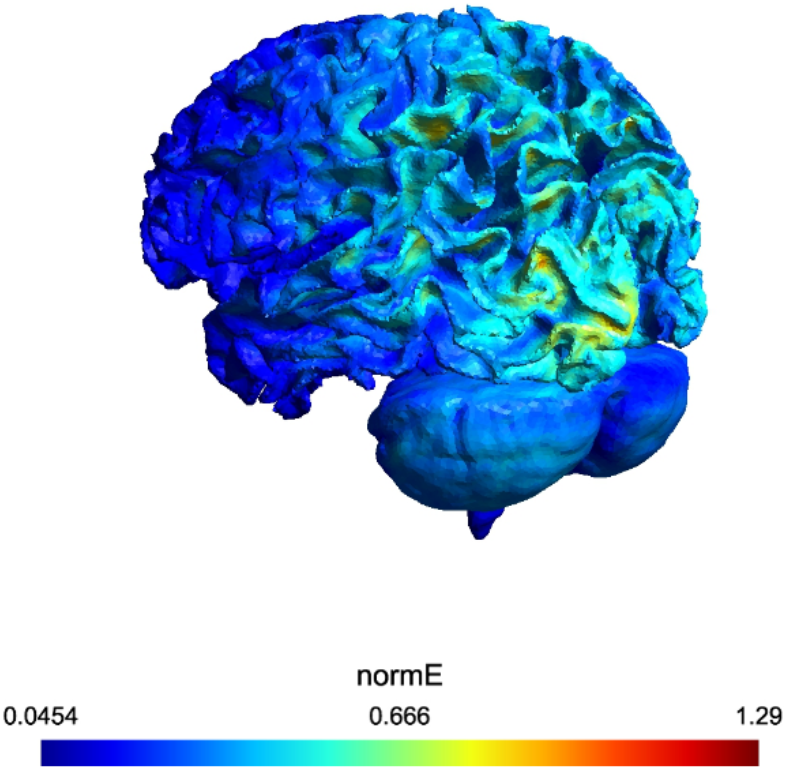
Modelled current flow in the brain. This image shows where the current flows in the brain. It does not provide any information on the frequency characteristics, therefore, we modelled this separately (see table 1).

## Discussion and conclusion

Our results demonstrate for the first time that adding noise to a visual stimulus during binocular rivalry can significantly reduce mixed percept durations. The same results were obtained when noise was added to the visual cortex directly with tRNS. These effects coincided with a significant reduction in the standard deviation of the mixed percept duration, suggesting less variability in the mixed dominance durations. This effect only occurred when the visual stimuli had a low contrast. These results were consistent with the predictions made by a computational binocular rivalry model (see fig. 2). Exclusive percept dominance durations in experiment 1, 2 and 3 were not affected by noise.

Adding noise to the visual stimuli (peripheral noise) resulted in the same behavioural results as adding noise to the brain directly with tRNS (central noise), that is, a significant reduction in mixed percept duration when the stimuli had a low contrast. No effect was observed when the stimuli were presented with a high contrast, suggesting that the stimulus contrast is a crucial parameter. Previously we have shown that noise added peripherally and centrally can both enhance performance on a contrast detection task (van der Groen & Wenderoth, 2016) for weak visual stimuli according to a SR-effect. Unpublished findings suggest that this is due to an increase in the quality of the signal representation in the brain. An important difference between these two noise application methods is the location in the nervous system where they can influence the visual processing for the first time. Peripheral noise likely influences visual processing at the receptor level, before the visual stimuli are processed by the brain. tRNS on the other hand influences visual processing at the cortical level. By applying tRNS during binocular rivalry we show that noise at a cortical level can causally influence mixed percept duration.

The processes underlying the occurrences of mixed percept during binocular rivalry are unclear, however, a possible mechanism involves changes in the strength of mutual inhibition between the two neuronal pools, each coding for a different percept (Hollins, 1980; Klink, Brascamp, Blake, & van Wezel, 2010). It is thought that stronger stimuli (stimuli with a higher signal-to-noise ratio) result in an increase in mutual inhibition, which leads to a reduction of the mixed percept duration (Hollins, 1980). Adding noise could have strengthened mutual inhibition due to an increase in the SNR of the stimulus representation in the brain. Related to this finding, a recent study demonstrated that alcohol intake enhances mixed percept duration (Cao, Zhuang, Kang, Hong, & King, 2016). It has been shown in cats that alcohol reduces the signal-to-noise ratio (i.e. increases noise) in primary visual cortex (Chen, Xia, Li, & Zhou, 2010). In our study we likely increased the SNR in visual cortex, resulting in a reduction of mixed percept duration. Another possible mechanism that effects mixed percept duration is a change in the balance between excitatory and inhibitory neural activity (Said, Egan, Minshew, Behrmann, & Heeger, 2013). An imbalance between cortical excitation and inhibition is an important factor in autism spectrum disorder (ASD) models (Robertson, Kravitz, Freyberg, Baron-Cohen, & Baker, 2013). However, it is still unknown whether lower biological noise levels are also involved in ASD (Davis & Plaisted-Grant, 2015). Interestingly, our finding that only mixed percepts are sensitive to a noise effect is supported by a study into binocular rivalry in ASD (Robertson et al., 2013). Robertson and colleagues did not find any difference between healthy controls and people with ASD on exclusive percept duration but an increase in mixed percept duration in people with ASD.

It is known that tRNS is able to enhance cortical excitability after 4 minutes of continuous stimulation (Chaieb, Paulus, & Antal, 2011), which results in a change in the excitation-inhibition balance. However, with our design it is unlikely that we changed cortical excitability because of the relatively short tRNS application durations. Moreover, after each tRNS block there was always a block without tRNS to reduce the likelihood of any after effects. Besides this, adding noise to the visual stimulus resulted in similar effects, and as far as we know cortical excitability cannot be modulated by peripheral noise. Therefore, our results are most likely explained by a SR-effect that modulates perceived stimulus contrast. In experiment 3 we did not find any effect of noise on rivalry dynamics. This is likely not because binocular rivalry is insensitive to a SR-effect, since it has been previously demonstrated that binocular rivalry is sensitive to a SR-mechanism (Kim, Grabowecky, & Suzuki, 2006). In contrast to our study, Kim and colleagues did not add noise to the rivalry process, but changed the strength of the driving signal by changing the contrast of the two images in counter phase. The idea is that SR will occur when the driving signal matches biological noise levels. The reason that there is no noise effect on the dominance duration of the exclusive percepts in our study could be because the noise added in the current study might not be optimal to introduce a SR-effect in a bistable system. The noise properties we applied were based on the findings of our previous study (van der Groen & Wenderoth, 2016) where we demonstrated that tRNS and peripheral noise are able to enhance performance on a contrast detection task.

In order to determine whether tRNS induced electrical noise reached the cortex we used simulations based on a spherical head model (Truong et al., 2014). The modelling results demonstrated that our stimulation induced an electrical field around 2 V/m and that all frequencies were transmitted to the brain. With tRNS a mix of alternating currents (AC) is applied to the cortex. It has been calculated that an electrical field strength of 1 V/m of 100 Hz AC can polarize a neuron by only 50μV (Deans, Powell, & Jefferys, 2007). This estimated field strength is too small to directly depolarize neurons. However, it is thought that this signal can be amplified by stimulating more neurons simultaneously (Francis, Gluckman, & Schiff, 2003). This small depolarization may be enough to elicit action potentials (APs) in neurons that are close to threshold, which results in a weak stimulus reaching the threshold AP generation earlier.

Although it is still debated where in the brain binocular rivalry is resolved, we targeted the visual cortex. Our current spread modelling results (fig. 4) demonstrate that our setup was suitable for stimulation of the visual cortex. The reason we targeted the visual cortex is because neuronal activity in primary visual cortex (V1) is linked to the subjective percept during binocular rivalry (Lee, Blake, & Heeger, 2007; Leopold & Logothetis, 1996; Polonsky, Blake, Braun, & Heeger, 2000). Changes in higher order brain areas, like frontoparietal activity, are thought to be the result of perceptual alternations rather than the cause of the perceptual alternations (Knapen, Brascamp, Pearson, van Ee, & Blake, 2011). However, this is still an area of debate as there is also evidence that frontoparietal activity might play a causal role (Carmel, Walsh, Lavie, & Rees, 2010; Kanai, Carmel, Bahrami, & Rees, 2011; Zaretskaya, Thielscher, Logothetis, & Bartels, 2010). Evidence that mixed percepts may be more influenced by low-level visual processes rather than higher order processes (Antinori, Carter, & Smillie, 2017; Blake, Oshea, & Mueller, 1992) supports the interpretation that our effect is driven by stimulation of these low-level visual cortices.

In conclusion, our results are the first to demonstrate that mixed percept durations can be altered by applying noise to the visual cortex directly with tRNS, likely due to an enhancement of stimulus contrast. Our results open up new ways of manipulating noise levels within the brain and provide a better understanding of the role noise plays in the brain when it is in a dynamical state.

